# The *C. elegans* heterochronic gene *lin-28* coordinates the timing of hypodermal and somatic gonadal programs for hermaphrodite reproductive system morphogenesis

**DOI:** 10.1101/265314

**Authors:** Sungwook Choi, Victor Ambros

## Abstract

*C. elegans* heterochronic genes determine the timing of expression of specific cell fates in particular stages of developing larva. However, their broader roles in coordinating developmental events across diverse tissues has been less well investigated. Here, we show that loss of *lin-28*, a central heterochronic regulator of hypodermal development, causes reduced fertility associated with abnormal somatic gonad morphology. In particular, the abnormal spermatheca-uterine valve morphology of *lin-28(lf)* hermaphrodites trap embryos in the spermatheca, which disrupts ovulation and causes embryonic lethality. The same genes that act downstream of *lin-28* in the regulation of hypodermal developmental timing also act downstream of *lin-28* in somatic gonad morphogenesis and fertility. Importantly, we find that hypodermal expression, but not somatic gonadal expression, of *lin-28* is sufficient for restoring normal somatic gonad morphology in *lin-28(lf)* mutants. We propose that the abnormal somatic gonad morphogenesis of *lin-28(lf)* hermaphrodites results from temporal discoordination between the accelerated hypodermal development and normally timed somatic gonad development. Thus, our findings exemplify how a cell-intrinsic developmental timing program can also control cell non-autonomous signaling critical for proper development of other interacting tissues.

## Introduction

The term “heterochrony” refers to modes of developmental alterations of an organism in which genetic changes lead to either accelerated or delayed development of certain body parts relative to others, in the context of evolution (Keyte and Smith, 2012; Klingenberg, 1998), or in experimental organisms (Ambros and Horvitz, 1984; Ambros and Horvitz, 1987). In the evolutionary context, heterochrony can cause characteristic morphological differences between an organism and its ancestor (Tompkins, 1978). Genetic mutations causing heterochrony in the nematode *Caenorhabditis elegans* identified a gene regulatory network, the ‘heterochronic pathway’, that governs the relative timing of developmental events during the four larval stages (L1~L4), particularly in the hypodermis. Loss-of-function (lf) mutations of the heterochronic genes *lin-14* or *lin-28* result in precocious development via the skipping of hypodermal cell fates specific to one or several larval stages (Ambros and Horvitz, 1984; Ambros and Horvitz, 1987). In contrast, *lin-4(lf)* and *let-7(lf)* mutations prevent the normal progression of certain stage-specific cell fates, leading to abnormal repetition of larval stages (Ambros and Horvitz, 1984; Reinhart et al., 2000).

*lin-4* and *let-7* encode microRNAs that post-transcriptionally control the expression of the protein products of certain other heterochronic genes (Lee et al., 1993; Reinhart et al., 2000). The *lin-4* microRNA binds to the 3’ untranslated region (UTR) of the *lin-14* and *lin-28* mRNAs to inhibit their translation, and *let-7* negatively regulates other heterochronic genes, including *hbl-1* and *lin-41* (Abrahante et al., 2003; Slack et al., 2000).

The *lin-28* gene encodes a conserved RNA-binding protein containing one cold shock domain and two zinc finger domains. Notably, the mammalian homolog of *lin-28* is implicated in diverse biological processes, including tumorigenesis, pluripotency, and metabolism (Asara et al., 2016; Piskounova et al., 2011; Yu et al., 2007; Zhu et al., 2011). In *C. elegans,* expression of LIN-28 protein is highest in the late embryo and L1 stages and decrease from the L2 stage onwards. LIN-28 is primarily expressed in the hypodermis, neurons, and muscle. *lin-28(lf)* mutants skip L2-specific hypodermal cell fates, resulting in precocious expression of L3, L4 and adult hypodermal fates (Moss et al., 1997; Seggerson et al., 2002). Thus, in the wild type, *lin-28* functions in early larval stages of the worm to specify the proper timing of L2-to-L3 cell fates transition. *lin-28* does this by positively regulating expression of the HBL-1 transcription factor, and by preventing premature expression of mature *let-7* microRNA (Abbott et al., 2005; Vadla et al., 2012; Van Wynsberghe et al., 2011). The roles of LIN-28 in the regulation of *let-7* are conserved in many other species including mammals (Mayr et al., 2012; Stratoulias et al., 2014; Viswanathan et al., 2008).

Here, we investigated the role of *lin-28* in maintaining fertility of *C. elegans* hermaphrodites. The hermaphrodite gonad consists of germ cells as well as non-germ cell tissues that are essential for gonadal function and fertility. While germ cells originate from proliferation of the Z2 and Z3 germline precursor cells of L1 larvae, all somatic gonadal tissues are derived from Z1 and Z4 cells, located within the gonadal primordium next to the two germ line precursors (Kimble and Hirsh, 1979). Somatic gonad development occurs during the L1-L4 larval stages, and is characterized by stage specific patterns of cell division, morphogenetic events, differentiation of cells into tissues with specific functions, including the anchor cell (AC), distal tip cells (DTCs), the gonadal sheath cells, the spermatheca, the spermathecal-uterine valve (Sp-Ut valve), the uterus, and the uterine seam cells (utse). Mature oocytes are released during ovulation to the spermatheca for fertilization. Post-fertilization, the embryo exits into the uterus where embryogenesis begins. Both ovulation and spermathecal exit are important reproductive processes in *C. elegans* hermaphrodites, and mutants defective in these processes have been shown to exhibit reduced fertility (Iwasaki et al., 1996; Kariya et al., 2004; Kovacevic and Cram, 2010).

Our results show that certain aspects of somatic gonad development are abnormal in *lin-28(lf)* hermaphrodites, reflected by abnormal morphology of the uterus, uterine seam, and Sp-Ut valve. These morphological defects, particularly the abnormal Sp-Ut, dramatically limit *lin-28(lf)* fertility. Our results further indicate that the normal development of the somatic gonad relies on temporal coordination of hypodermal developmental events with somatic gonadal events, and that *lin-28* acts in the hypodermis to specify a schedule of hypodermal events that is properly coordinated with a corresponding schedule of somatic gonadal developmental events. We demonstrate that the hypodermal function of *lin-28* is sufficient to regulate somatic gonad development non-autonomously, consistent with a role for *lin-28,* and downstream heterochronic genes, in controlling the hypodermal components of critical developmental signaling between the gonad and hypodermis.

## Results

### *lin-28(lf)* mutants exhibit defects in embryo production and embryo viability

*lin-28(n719)* hermaphrodites produced dramatically fewer larval progeny than wild-type hermaphrodites, with the reduced fertility most extreme at 25°C (Fig 1A). *lin-28(n719)* mutants are unable to lay eggs, due to their precocious vulva formation, which results in failure of proper vulva morphogenesis (Euling and Ambros, 1996). Like other egg-laying defective mutants, *lin-28(lf)* hermaphrodites contain their entire brood of embryos trapped inside the limited space of the somatic gonad. To test whether the reduced number of progeny of *lin-28(lf)* hermaphrodites could be simply the result of their egg-laying defects, we compared the number of progeny of *lin-28(n719)* hermaphrodites with that of *lin-2(e1309). lin-2(e1309)* mutants exhibit defective egg-laying due to their vulvaless phenotype, which results from cell lineage defects not related to developmental timing (Hoskins et al., 1996). *lin-28(n719)* mutants produce substantially fewer progeny than *lin-2(e1309)* mutants (Fig 1A), suggesting the reduced brood size of *lin-28(n719)* animals is caused by a combination of factors, such as reduced embryo production or elevated embryonic lethality, compared to *lin-2(e1309).* To test these two possibilities, we first dissected mature, gravid hermaphrodites to count the embryos retained inside. *lin-28(n719)* mutants had fewer embryos than *lin-2(e1039)* mutants in gravid adults (Fig 1B), indicating that *lin-28(n719)* mutants are defective in embryo production. We also harvested embryos from dissected gravid adults and counted the number that hatched and developed into larva. Approximately 70% of *lin-28(n719)* embryos failed to develop, whereas essentially all of the *lin-2(e1309)* embryos were viable (Fig 1C). These results suggest that both reduced embryo production and embryonic lethality contribute to the fertility defects in *lin-28(n719)* mutants. *lin-28(n719)* mutants also showed the same fertility defects at 20°C, caused by reduced embryo production and embryonic lethality, but at a somewhat reduced penetrance compared to 25°C (Fig S1). We conducted all our subsequent experiments at 25°C, where the fertility defects are prominent.

**Figure 1.**
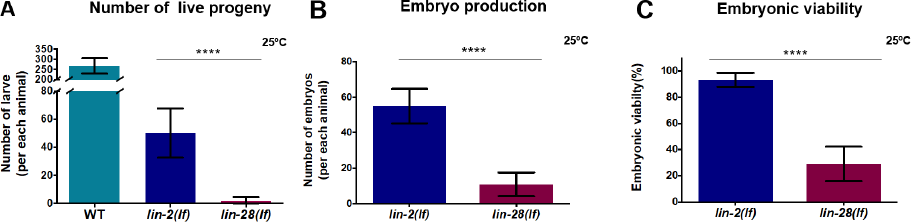
*lin-28(lf)* hermaphrodites have reduced brood size and exhibit defects both in embryo production and embryonic viability. (A) Number of live progeny per hermaphrodite for wild type (strain N2) (n=5) and for *lin-2(e1309)* (n=35) and *lin-28(n719)* mutants (n=104) at 25°C. *lin-2(e1309)* mutants are used as egg-laying defective controls; both *lin-2(e1309)* and *lin-28(n719)* hermaphrodites are unable to lay eggs. (B) Number of embryos per hermaphrodite for *lin-28(n719)* (n=12) and *lin-2(e1309)* (n=10) mutants at 25°C (dissected from gravid adults 60 hr after initiation of synchronized larval development by feeding starved L1s). (C) Viability of mutant embryos dissected from *lin-28(n719)* and *lin-2(e1309)* hermaphrodites at 25°C. % viability = 100×(viable hatched larvae/total embryos). (Number of animlals≥15 per each assay; number of independent replicate assays = 11 for *lin-28(n719),* 4 for *lin-2(e1309))* (A-C: Data are shown as mean ± SD, unpaired t-test, ****p<0.0001)

### Defects in ovulation and the spermathecal exit cause reduced embryo production in *lin-28(lf)* mutants

To investigate the cause of reduced embryo production in *lin-28(n719)* mutants, we checked whether ovulation (the passage of mature oocytes into the spermatheca, where fertilization occurs) and spermathecal exit (the passage of fertilized embryos from the spermatheca into the uterus) proceed normally in the mutants. In the wild type, fertilized embryo exit the spermatheca through the Sp-Ut valve to the uterus (McCarter et al., 1999). We examined the spermatheca of *lin-28(n719)* mutants using the expression of a spermatheca reporter, *fkh-6::GFP* (Chang et al., 2004). Approximately 80% of *lin-28(n719)* mutants contained embryos in their spermathecae, whereas less than 10% of wild-type spermathecae contained embryos (Fig 2A,B). Unlike in the wild type, many embryos in *lin-28(n719)* hermaphrodites had undergone multiple rounds of cell division inside the spermathecae. This suggests that *lin-28(n719)* mutants have defects in the process of spermathecal exit. We used time-lapse video microscopy to monitor spermathecal exit (Video S1,2). In wild type hermaphrodites, the process of ovulation, fertilization, and spermathecal exit of an individual embryo happen within less than 10 minutes (McCarter et al., 1999). In *lin-28(n719* mutants however, an embryo was trapped in the spermatheca and unable to exit into the uterus for 40 minutes (Video S2).

**Figure 2.**
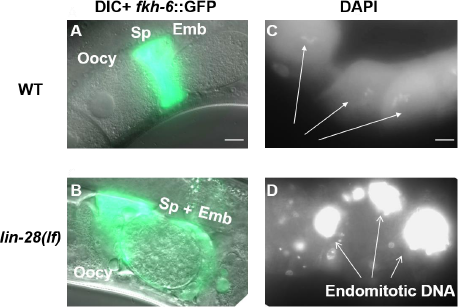
*lin-28(lf)* hermaphrodites have defects in exit of embryos from the spermatheca, and defects in ovulation. (A, B) Spermatheca labeled by *fkh-6:GFP* of representative wild-type and *lin-28(n719)* adult hermaphrodites. (A) In the wild type, oocytes (Oocy) pass into the spermatheca (Sp), where they are fertilized, and rapidly exit as a one-cell embryo (Emb), and so most spermathecae are not observed to contain an embryo. (B) In this *lin-28(n719)* hermaphrodite, an embryo (around ~150 cells) was trapped in the spermatheca. (C, D) DAPI staining of oocytes in wild-type (C) and *lin-28(n719)* hermaphrodites. (C) In the wild type, individual oocytes (Oocy) contained a haploid complement of condensed chromosomes (arrows). (D) In this *lin-28(n719)* mutant, endomitotic DNA was evidenced by an excessively bright DAPI signal, a characteristic of ovulation defective mutants. (Scale bar = 10 µm in this paper, except where noted).

Next, we addressed whether the process of ovulation is also defective in *lin-28(n719)* mutants. Endomitotically replicating DNA (Emo) oocytes is a characteristic of many ovulation mutants (Iwasaki et al., 1996), where oocytes undergo several DNA replications without ovulation and fertilization. Gonadal DAPI staining revealed that some *lin-28(n719)* hermaphrodites contain endomitotic oocytes (Fig 2D). We speculate that the defective ovulation in *lin-28(n719)* mutants may result from impairment of the spermathecal exit process, wherein the presence of fertilized embryos trapped within the limited spermathecal space would prevent entry of mature oocytes. We conclude that the poor fertility of *lin-28(n719)* mutants is the consequence of embryos becoming trapped in the spermathecae, resulting in inhibition of the ovulation process.

### Sp-Ut valve morphogenesis, uterus lumen formation, and utse migration are abnormal in *lin-28(lf)* mutants

We wondered whether *lin-28(n719)* mutants are defective in passage of embryos through the spermatheca on account of defects in proliferation of spermathecal progenitor cells. Spermathecae in *C. elegans* hermaphrodites consist of two rows of 12 cells forming a long tube structure at the young adult stage (Gissendanner et al., 2008; Kimble and Hirsh, 1979). Long spermathecal tube structures, labeled by *fkh-6:GFP,* are also observed in the *lin-28(n719)* mutants at the 4^th^ stage (Fig 3B), which corresponds to L4 and adult stage in wild type (Fig 6C). Wild-type spermathecae become constricted horizontally as somatic gonadal tissues continue morphogenesis before the first ovulation. *lin-28(n719)* spermathecae exhibited a similar constricted and extended morphology as in the wild type (Fig 3A,B). Overall, we did not detect appreciable differences in spermathecal morphology between wild-type animals and *lin-28(n719)* mutants. The Sp-Ut valve connects the spermatheca to the uterus in wild type hermaphrodites. The mature Sp-Ut valve consists of the toroidal syncytium and the core cell syncytium. We examined the structure of the Sp-Ut valve in *lin-28(n719)* mutants according to the expression of *cog-1:GFP,* which is expressed in the Sp-Ut valve core cell syncytium from the late L3 or early L4 stage (Palmer et al., 2002). In wild type animals, the core cell syncytium stretches during the L4 stage, and in young adults, the Sp-Ut valve core forms a dumbbell-like structure with one end in the spermatheca and the other end in the uterus (Fig 3C). Although we observed apparently normal expression of *cog-1:GFP* in the Sp-Ut valve region of *lin-28(n719)* mutants in late 3^rd^ larval stage, the stretching of a core cell syncytium did not occur in the mutants and the Sp-Ut valve core remained as a single lobe structure in old 4^th^ stage and later (Fig 3D). This observation indicates that the Sp-Ut valve morphology is abnormal in *lin-28(n719)* mutants, suggesting the connection between the spermatheca and the uterus is also disrupted in the mutants. We conclude that the aberrant connection between the uterus and the spermatheca induced by the atypical morphology of the Sp-Ut valve causes the spermathecal exit defect in *lin-28(lf)* mutants.

**Figure 3.**
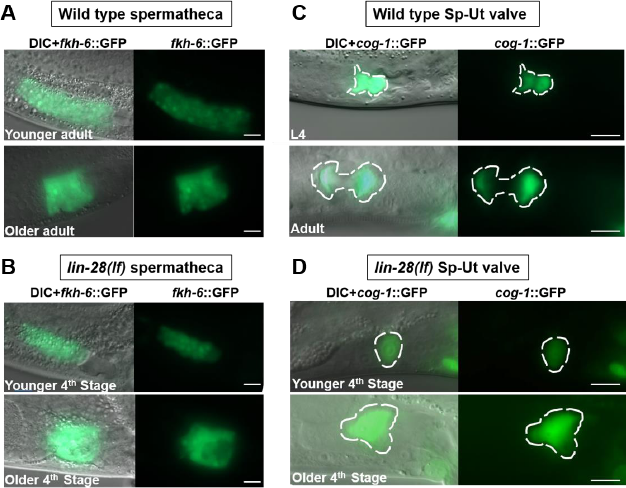
*lin-28(lf)* mutants show essentially normal spermathecal primordium structure, but display defects in spermathecal-uterine (Sp-Ut) valve morphology. Spermathecal primordium visualized by *fkh-6:GFP* expression (A, B) and Sp-Ut valve core structure visualized by cog-1:GFP expression (C, D) in wild-type and *lin-28(n719)* hermaphrodites at successive times in the advancement towards first ovulation. In each panel, the upper images are of hermaphrodites at the L4 or early young adult stage, and the lower images are of hermaphrodites somewhat later in development, just before the time of first ovulation. *(lin-28(n719)* hermaphrodites skipped one larval stage; therefore, the “4th stage” corresponds to L4 and adult stage in wildtype [See Fig 6].) (A, B) In both wild type and *lin-28(n719)* mutants, similar tube-shaped spermathecal primordia were detected, which contracted horizontally to form similar sac-like structures in older adults (A, B lower panels). (C, D) Sp-Ut valve core structure (Sp-Ut, dashed oval) labeled by cog-1:GFP at successive stages in L4-adult developmental progression of a wild type and *lin-28(n719)* hermaphrodite. (C) In wild type, the Sp-Ut valve core stretches to form a fully developed “dumbbell” structure, with one side residing in the spermatheca and the other side in the uterus. The distance between each side shown here was ~10 µm. (D) In *lin-28(n719)* mutants, the Sp-Ut valve core does not stretch and remained as “single lobe” structure, indicating an abnormal connection between the spermatheca and uterus in the mutants

To assess whether the abnormal Sp-Ut valve morphology of *lin-28(n719)* could reflect somatic gonadal cell lineage defects analogous to their precocious hypodermal cell lineages (Ambros and Horvitz, 1984), we investigated whether the Sp-Ut valve syncytium in the mutant contains a normal number of nuclei as in the wild-type. The Sp-Ut valve is derived from daughter cells of the dorsal uterus lineage, and the valve core syncytium is comprised of a fusion of two cells (Kimble and Hirsh, 1979). We used confocal microscopy to count the number of nuclei in the Sp-Ut valve core region based on labeling by *cog-1:GFP.* Two nuclei were present in Sp-Ut core cell of both wild-type and *lin-28(n719)* mutants, suggesting the morphological defect is not caused by abnormal cell division in the lineage generating the Sp-Ut valve core (Fig S2A,B).

Additional somatic gonadal defects were also detected in *lin-28(n719)* mutants. In wild-type animals, the uterus lumen forms between the dorsal and ventral uterus during the L4 stage when uterus toroidal cells fuse to generate a syncytium (Newman et al., 1996). In the *lin-28(n719)* mutants, several small lumens were observed in the uterus region, but completely connected lumens were not detected in most of the mutants during the 4^th^ stages (Fig S3B). Some *lin-28(n719)* mutants had connected lumens, but the length of the lumen was shorter and the overall lumen shape was rounder than in wild-type animals (Fig S3C).

The utse represents a component of the hermaphrodite somatic gonad that mediates structural attachment between the hypodermis and uterus (Newman and Sternberg, 1996). The utse syncytium forms in early L4 stages and extends laterally during early to mid L4. The utse connects the uterus to the seam cells laterally and also to uv1 and vulva cells. *egl-13:GFP* is expressed in the nuclei of π cell lineage, whose products includes the utse, from late L3 to L4 stages (Ghosh and Sternberg, 2014). Utse labeled by *egl-13:GFP* migrated laterally in wild-type hermaphrodites (Fig S4A,B). However, the utse nuclei did not migrate during the 4^th^ stages in *lin-28(n719)* mutants, although *egl-13:GFP(+)* cells were detected in the utse region (Fig S4C,D).

*lin-28(n719);lin-2(e1309)* double mutants, which lack vulva formation, have the same somatic gonadal defects as *lin-28(n719)* mutants, indicating these defects are not indirect consequences of the abnormal vulva morphology in *lin-28(n719)* mutants (Fig S5).

Defects in uterus lumen formation and utse migration in *lin-28(n719)* mutants, together with Sp-Ut valve morphological defects, suggest that *lin-28* activity is required for multiple aspects of proper hermaphrodite somatic gonad development.

### Eggshell integrity is compromised in *lin-28(lf)* mutant embryos

To understand the nature of embryo lethality of *lin-28(n719)* mutants, we dissected and observed the embryos produced by the mutants. The *lin-28(n719)* embryos displayed abnormal, irregular shapes (Fig 4A), reminiscent of the misshapen phenotypes of many egg-shell mutants (Johnston et al., 2006; Maruyama et al., 2007; Zhang et al., 2005). The *C. elegans* eggshell is formed rapidly after fertilization in the spermatheca and consists of chitin, lipid, and other structural proteins protecting the embryo. Eggshell abnormalities often results in embryo lethality (Johnston et al., 2010). We first confirmed whether *lin-28(n719)* embryos have an eggshell. Chitin-binding domain-protein 1 (CBD-1) is a component of the eggshell cortex, and thus, CBD-1::mcherry is expressed all around the wild-type embryos. Despite the differences in embryo shapes, CBD-1::mcherry expression was observed along the periphery of *lin-28(n719)* embryos, implying that the eggshell forms in the mutant embryos (Fig 4B). However, *lin-28(n719)* embryos were abnormally permeable to the lipophilic dye FM 4-64. The wild-type eggshell functions as a barrier between the embryo and outside environment, which prevents the lipophilic dye from infiltrating the embryo (Johnston et al., 2006). In our study, about 50% of *lin-28(n719)* embryos were permeable for FM4-64, while around 10% of *lin-2(e1309)* embryos were permeable (Fig 4C). This finding indicates that eggshell integrity is compromised in *lin-28(n719)* mutants. Interestingly, embryos of *fln-1(tm545)* mutants, which have defective spermathecal exit (Kovacevic and Cram, 2010), are also more permeable to FM4-64 than *lin-2(e1039)* embryos, implying a potential association of eggshell permeability with the spermathecal exit phenotype (Fig 4C). A delay in the spermathecal exit might decrease the physical integrity of eggshell in the *lin-28(lf)* mutants (see Discussion). In addition, *lin-28(n719)* embryos are more sensitive to osmotic stresses. Wild-type embryos maintained their oval shape upon exposure to 0, 150, and 300 mM KCl, whereas *lin-28(n719)* embryos could not maintain the shape and swelled upon treatment with 0 or 150mM of KCl (Fig 4D). Therefore, *lin-28(n719)* eggshells are not able to function as a permeable and osmotic barrier, and thereby contribute to the embryonic lethality.

**Figure 4.**
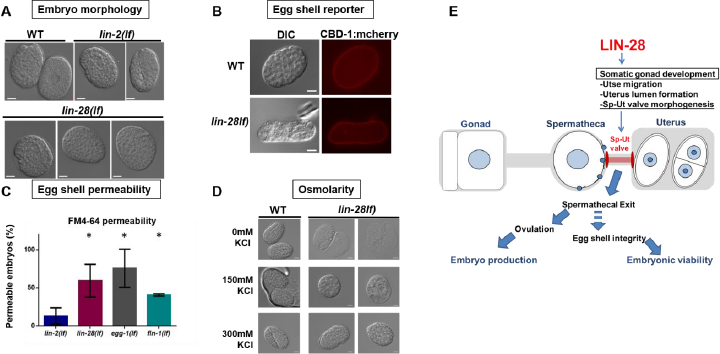
*lin-28(lf)* embryos are misshapen and are defective in egg shell integrity. (A) DIC images of wild-type, *lin-2(e1309)* and *lin-28(n719)* embryos from dissected adult animals. Wild type and *lin-2(e1309)* mutants produce oval embryos, but *lin-28(n719)* embryos exhibit irregular shapes. (B) *cbd-1::mCherry* expression indicates that egg shells were present in *lin-28(n719)* embryos despite their misshapen morphology (lower panels). (C) Egg shell permeability of embryos produced by *lin-2(e1309), lin-28(n719), egg-1(tm1071),* and *fln-1(tm545)* hermaphrodites. Egg shell mutant *egg-1(tm1071)* served as a control. Embryos from *lin-28(n719), fln-1(tm545),* and *egg-1(tm1071)* were more permeable to the lipophilic dye FM 4-64 than *lin-2(e1309)* embryos. Like *lin-28(n719), fln-1(tm545)* hermaphrodites exhibit defects in spermathecal exit. Permeability was calculated as the percentage of permeable embryos/total embryos from dissected adult animals (Number of animlals≥15 per each assay; number of independent replicate assays = 3 for each strain, Data are shown as mean ± SD. unpaired t-test compared to *lin-2(e1309),* *p<0.05). (D) Morphology of embryos under different osmotic conditions. *lin-28(n719)* embryos were more sensitive than wild type to low salt conditions, indicating a lack of protection from osmotic stress. (E) Model for physiological causes of fertility defects in *lin-28(lf)* mutants. *lin-28(lf)* animals have abnormal Sp-Ut valve structure, which leads to defects in spermathecal exit and ovulation, and hence a reduced embryo production. In addition, retention of embryos in the spermatheca compromises egg shell integrity, which causes embryonic lethality.

In summary, *lin-28* is required for normal somatic gonad development, including uterine lumen formation, utse morphogenesis, and proper Sp-Ut valve formation. The abnormal Sp-Ut valve structure in *lin-28(n719)* mutants causes defects in spermathecal exit and ovulation, resulting in impaired embryo production. In addition, we hypothesized that the impaired eggshell integrity in mutants is caused by a spermathecal exit defect that leads to embryonic lethality (Fig 4E).

### Embryonic lethality of *lin-28(n719)* mutant is rescued by maternal expression of *lin-28*

We speculated that if the embryonic lethality of the *lin-28(n719)* mutant is due to defects in maternal somatic gonadal morphology, then maternal expression of *lin-28* should rescue the embryonic lethality of these mutants. To test this, we crossed *lin-28(n719)* mutants with wild-type males to obtain heterozygous mutants *(lin-28(n719)/+).* These heterozygotes appeared to exhibit essentially normal fertility, in terms of overall numbers of viable progeny. We then assessed the viability of *lin-28(n719)* homozygous self-progeny from these *lin-28(n719)/+* hermaphrodites. Viable *lin-28(n719)* homozygotes were identified by their characteristic egg-laying defective phenotype as adults. Amongst the self-progeny of heterozygous *(lin-28(n719)/+)* hermaphrodites, we observed a ratio of egg-laying capable progeny (+/+ or *lin-28(n719)/+)* to egg-laying defective progeny *(lin-28(n719)/lin-28(n719))* of 2.89(±0.2):1 which is very close to the expected 3:1 ratio for complete maternal rescue of embryonic lethality (Table 1).

**Table 1.** Viability of embryos produced by *lin-28(n719)* heterozygous hermaphrodites

| Maternal Genotype (P0) | Progeny Genotype (F1) | Maternal or Zygote LIN-28 products | Egg laying Phenotype (F1) | Expected Genotype ratio (Zygotic effect) | Expected Phenotype ratio (Zygotic effect) | Expected Genotype ratio (Maternal Effect) | Expected Phenotype ratio (Maternal Effect) | Actual Ratio (by egg laying phenotype) |
| --- | --- | --- | --- | --- | --- | --- | --- | --- |
| *lin-28(+)*<br>*lin-28(-)* | *lin-28(+)*<br>*lin-28(+)* | m+/z- | Egg laying | 1 | 10.3 | 1 | 3 | 2.89(±0.2)** |
|  | *lin-28(+)*<br>*lin-28(-)* | m+/z- | Egg laying | 2 |  | 2 |  |  |
|  | *lin-28(-)*<br>*lin-28(-)* | m-/z- | Egg laying defective | 0.29* | 1 | 1 | 1 | 1 |

\*Average embryo viability of *lin-28(lf)* mutants is 71%
\*\* Calculated by progeny from 8 individual *lin-28(n719)* heterozygotes

In this experiment, if *lin-28(n719)* embryonic lethality were solely due to a zygotic role of *lin-28* in embryos, the expected ratio of egg-laying progeny (+/+ or *lin-28(n719)/+)* to egg-laying defective progeny *(lin-28(n719)/lin-28(n719))* would have been 10:1 (Table 1). Instead, we observed a ratio of egg-laying progeny to egg-laying defective progeny very close to the 3:1 Mendelian ratio, indicating that *lin-28(n719)* embryonic lethality reflects a requirement for *lin-28* activity in the mother to enable proper development of the somatic gonad, which is required for the production of viable embryos.

### Genes downstream of *lin-28* in developmental timing regulation also function downstream of *lin-28* in somatic gonad morphogenesis and fertility

Genetic interactions of *lin-28* with other heterochronic genes in *C. elegans* have been described in previous studies (Ambros, 2011; Resnick et al., 2010). *hbl-1* is positively regulated by *lin-28* and prevents precocious transition from L2 to L3 fates in hypodermal cell lineages. *lin-46* inhibits the function of *hbl-1,* and thus, loss of *lin-46* suppresses the hypodermal developmental defects in *lin-28(n719)* mutants. In addition, *lin-28* prevents precocious biogenesis of mature *let-7* microRNAs. *hbl-1* is also negatively regulated by *let-7* in the L4 stage. *lin-29* acts downstream of *let-7* and *hbl-1* to positively regulate the transition from L4 fates to adult fates in the hypodermis (see Fig 5E, Developmental Timing).

**Figure 5.**
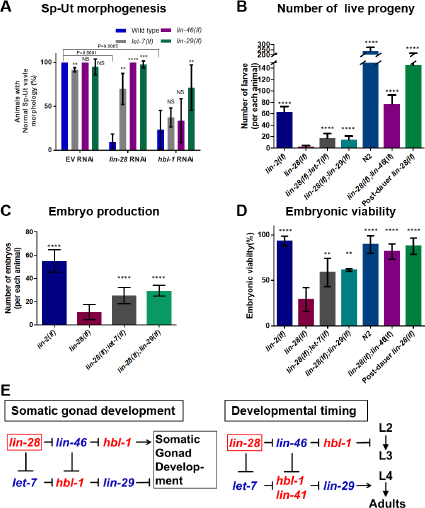
Genetic epistasis analysis of lin-28 and other heterochronic genes for effects on Sp-Ut valve morphogenesis and fertility. (A)The percentage of animals with normal Sp-Ut valve morphology, visualized by *cog-1*: GFP expression in wild type, *let-7(mn112), lin-46(ma164)* and *lin-29(n836)* mutants, treated with control RNAi (EV, empty vector strain L4440), lin-28(RNAi) or *hbl-1(RNAi).* Wild type, *lin-46(ma164),* and *lin-29(n836)* mutants treated with L4440 empty vector RNAi rarely showed Sp-Ut valve defects. *let-7(mn112)* mutants with control RNAi showed less than 10% of Sp-Ut valve defects. lin-28(RNAi) treatment of wild type led to ~95% Sp-Ut valve defects, an effect that was partially or fully suppressed by *let-7(mn112), lin-46(ma164)* and *lin-29(n836).* hbl-1(RNAi) treatment of wild type also led to Sp-Ut valve defects that were rarely suppressed by *let-7(mn112)* and *lin-46(ma164)* and moderately suppressed (~70%) by *lin-29(n836).* (Number of animlals ≥12 per each assay; number of independent replicate assays = 5 for WT with *hbl-1* RNAi, 4 for *lin-46(ma164)* or *lin-29(n836)* with *hbl-1* RNAi, 3 for all other assays, Data are shown as mean ± SD, unpaired t-test, p-values denoted as asterisks or “NS” were calculated by each mutant compared to wild type in each RNAi set. NS; not significant, **p<0.01, ***p<0.001, ****p<0.0001) (B-D) Fertility phenotypes of *lin-2(e1309), lin-28(n719), lin-28(n719);let-7(mn112), lin-28(n719);lin-29(n836), lin-46(ma164);lin-28(n719)* mutants, post-dauer *lin-28(n719),* and wild type. (B)Total number of viable larva progeny of *lin-2(e1309)* (n=16), *lin-28(n719)* (n=64), *lin-28(n719);let-7(mn112)* (n=62), *lin-28(n719);lin-29(n836)* (n=41), *lin-46(ma164);lin-28(n719)* (n=17) mutants, post-dauer *lin-28(n719)* (n=9), and wild type (n=4). (C) Embryo production of *lin-2(e1309)* (n=10), *lin-28(n719)* (n=12), *lin-28(n719);let-7(mn112)* (n=24), *lin-28(n719);lin-29(n836)* (n=15) mutants.(D) Embryo viability of *lin-2(e1309), lin-28(n719), lin-28(n719);let-7(mn112), lin-28(n719);lin-29(n836), lin-46(ma164);lin-28(n719)* mutants, post-dauer *lin-28(n719),* and wild type. (Number of animlals≥15 per each assay; number of independent replícate assays = 11 for *lin-28(n719),* 4 for *lin-2(e1309),* 5 for *lin-28(n719);let-7(mn112),* and 3 for all other strains.) The reduction in the number of progeny of *lin-28(n719)* mutants is increased by loss of *let-7, lin-29* or *lin-46.* (B). Both embryo production (C) and embryo viability (D) are improved by loss of *let-7, lin-29* or *lin-46.* Total number of progeny and embryonic viability of *lin-28(lf)* mutants are also improved by post-dauer development (B,D). (B-D: Data are shown as mean ± SD, unpaired t-test compared to *lin-28(lf),* NS; not significant, **p<0.01,***p<0.001,****p<0.0001.) (E) The genetic regulatory pathway model for somatic gonadal morphogenesis, derived from the results of epistasis experiments presented in Fig 5A-D, is highly similar to the model for temporal regulation of hypodermal cell fates derived from previous studies (Ambros, 2011; Resnik & Rougvie 2010, See Discussion for *lin-41* involvement in the somatic gonad morphogenesis).

We conducted RNAi knockdown experiments to determine whether these same heterochronic genes that act downstream of *lin-28* for hypodermal cell fate timing are also involved in Sp-Ut valve morphogenesis downstream of *lin-28.* After treatment of *cog-1:GFP* hermaphrodites with *lin-28* RNAi, 95% of the animals showed an abnormal Sp-Ut valve morphology similar to that in *lin-28(n719)* mutants. However, around 70% of *lin-28(RNAi)*-treated *let-7(mn112);cog-1:GFP* hermaphrodites showed a normal Sp-Ut valve. Also, genetic absence of *lin-46* or *lin-29* abolished the Sp-Ut valve defects caused by *lin-28(RNAi).* Thus, *lin-46(lf), lin-29(lf)* or *let-7(lf)* can fully or partially rescue the Sp-Ut valve defects in *lin-28(RNAi)* animals (Fig 5A), supporting the idea that these heterochronic genes act downstream of *lin-28* for somatic gonad development. hbl-1(RNAi) phenocopied the fertility phenotypes of *lin-28(n719)* mutants, including abnormal Sp-Ut valve core morphology (Fig S6A,B). Loss of *lin-46* or *let-7* function rarely affected the Sp-Ut valve morphological defect in *hbl-1*(RNAi) animals, indicating *hbl-1* is epistatic to these genes. Finally, around 70% of *lin-29(lf);cog-1:GFP* animals showed the wild-type Sp-Ut valve morphology when *hbl-1* function was compromised (Fig 5A).

In addition, a normal uterus lumen was observed in ~50% of *lin-28(n719);let-7(mn112)* mutants (Fig S3D). Most *lin-28(n719)]lin-29(n836)* and *lin-28(n719);lin-46(ma164)* mutants showed complete uterus lumen formation (Fig S3E,F). Utse migration defects of *lin-28(lf)* mutants were also partially suppressed by loss of either *let-7* or *lin-46* (Fig S4G-J).

Consistent with the genetic epistasis observed above for somatic gonadal morphogenesis, *lin-46(lf), let-7(lf)* and *lin-29(lf)* were also epistatic to *lin-28(lf)* for fertility. The total numbers of live progeny per animal produced by the *lin-28(n719);let-7(mn112)* and *lin-28(n719);lin-29(n836)* double mutants were significantly higher than that produced by the *lin-28(n719)* mutants (Fig 5B). Also, the total live progeny number for the *lin-28(n719);lin-46(ma164)* mutants was even greater than that from *lin-2(e1309)* mutants, because the loss of *lin-46* rescued the vulva and egg-laying defects of *lin-28(n719)* mutants. *lin-28(n719);let-7(mn112)* and *lin-28(n719);lin-29(n836)* mutants produced more embryos per animal than *lin-28(n719)* mutants (Fig 5C), and embryonic viability was similarly suppressed in these double mutants, compared to *lin-28(n719)* (Fig 5D).

Overall, our findings indicate that *let-7, lin-46, hbl-1,* and *lin-29* act downstream of *lin-28* for somatic gonad development in a network configuration essentially identical to that previously-described for the control of hypodermal cell fate timing by these same genes (Fig 5E). To further investigate the relationship between hypodermal developmental timing and somatic gonadal morphogenesis, we examined whether post-dauer development can rescue the somatic gonadal defects in *lin-28(n719)* mutants. A previous study showed that the hypodermal heterochronic developmental defects of *lin-28(n719)* mutants, (including precocious vulva, precocious adult cuticle, and altered seam cell number) are efficiently suppressed if the mutants develop through the dauer larva (an alternative, temporarily arrested, third larval stage) followed by “post-dauer” developmental stages (Euling and Ambros VR, 1996; Liu and Ambros, 1991). We observed that after post-dauer development, *lin-28(n719)* adults exhibited restored fertility and embryo viability, normal morphology of the Sp-Ut valve, uterus lumen, and normal utse migration (Fig 5B,5D, S6E, S3G, S4E, and S4F). This suppression of somatic gonadal defects in *lin-28(n719)* animals by post-dauer development supports the supposition that *lin-28* acts indirectly, via downstream genes and events, to mediate normal somatic gonad morphogenesis.

### Heterochronic development between hypodermal tissue and somatic gonad tissues in lin-28(lf) mutants

Based on the above observations indicating parallels between the control of hypodermal developmental timing and somatic gonad morphogenesis by *lin-28* mutants, we investigated whether the stage-specificity of somatic gonadal developmental events might be altered in *lin-28(n719)* hermaphrodites, in analogy to their precocious hypodermal development. As a marker to monitor the expression of stage-specific programs in the somatic gonad and hypodermis, we employed *cog-1:GFP* (Palmer et al., 2002), which is expressed in the wild type dorsal uterus and Sp-Ut valve core, beginning from late-L3/early-L4 stage (Fig 6A). *cog-1:GFP* is also expressed in the ventral hypodermis (vulval cell lineages) beginning in the mid L4 stage, which is after the onset of *cog-1:GFP* somatic gonad expression (Fig 6B). Thus in the wild type, the somatic gonadal expression of *cog-1:GFP* appears first (late-L3/early-L4 stage) followed later (mid L4) by vulval expression.

**Figure 6.**
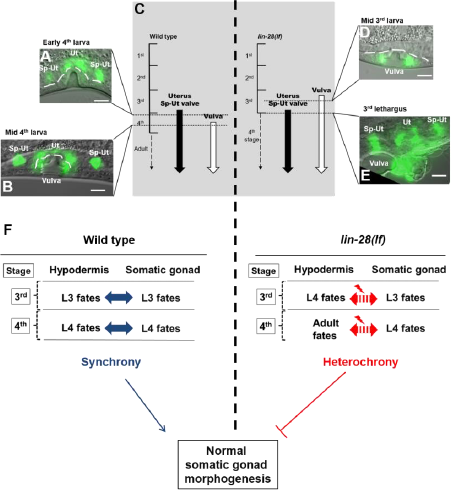
Heterochronic development of the hypodermis, relative to the somatic gonad, in *lin-28(lf)* hermaphrodites. (A,B,D,E) *cog-1:GFP* expression patterns in wild type and *lin-28(n719)* hermaphrodites at indicated developmental stages. The temporal order of the onset of cog-1:GFP expression in the somatic gonad and vulva is reversed in *lin-28(lf)* compared to the wild type. cog-1:GFP is expressed in both somatic gonadal tissues (uterus and Sp-Ut valve) and hypodermal tissue (vulva). The timing of expression in wild type and *lin-28(n719)* mutants is summarized in (C). In wild type, *cog-1:GFP* is expressed in somatic gonadal tissues at the late 3^rd^ larval stage (A) and in the vulva in the middle of the 4th larval stage (B). In *lin-28(n719)* mutants, vulva expression of *cog-1:GFP* occurs precociously in the middle of the L3 stages (D), while somatic gonadal expression of *cog-1:GFP* starts at the normal time, in the late 3^rd^ larval stage (E). Both vulval and somatic gonadal expression are detected at 3^rd^ lethargus (E). (F) Model for the importance of *lin-28* activity for somatic gonad development: In wild type animals, timing of hypodermis development and somatic gonad development are synchronized. In *lin-28(lf)* mutants, hypodermal development happens precociously while somatic gonad development does not, resulting in heterochronic development between two tissues. This heterochrony causes somatic gonad morphogenesis defects.

We examined whether the normal relative order of *cog-1:GFP* expression in the somatic gonad and vulva is altered in *lin-28(n719)* mutants. In *lin-28(n719)* hermaphrodites, *cog-1:GFP* expression in the vulva was observed precociously in the mid-3rd larval stage (Fig 6D), consistent with the previously-described precocious vulva cell divisions of *lin-28(lf)* mutants (Euling and Ambros, 1996). However, the onset of somatic gonadal expression of *cog-1:GFP* was normal, beginning from the late 3rd larval stage in both *lin-28(n719)* and in the wild type (Fig 6E). Thus, the expression of *cog-1:GFP* in the vulva precedes the expression in somatic gonads in *lin-28(n719)* mutants. These data suggest that the somatic gonad development, at least as reflected by *cog-1::GFP* expression, is not precocious in *lin-28(n719),* despite precocious development of the hypodermis. Therefore, developmental events in hypodermal tissues and somatic gonad tissues develop “heterochronicaNy” in *lin-28(n719)* mutants (Fig 6F). We hypothesize that this heterochronic development causes morphological defects of the somatic gonad, resulting in fertility defects.

### Hypodermal expression, but not somatic gonadal expression, of *lin-28* rescues normal somatic gonadal development and fertility in *lin-28(n719)* hermaphrodites

If the somatic gonadal morphological defects of *lin-28(n719)* mutants originate from discoordination between the timing of hypodermal and somatic gonadal development, suppressing the precocious hypodermal development in *lin-28(n719)* mutants should rescue such defects. We determined whether *lin-28* expression is necessary for hypodermal tissues to rescue the Sp-Ut morphological defects in the mutants. Using Mos1-mediated single copy insertion (MosSCI) transformation, we generated *lin-28:GFP::lin-28* 3’ UTR insertional strains driven by different Promoters: a *lin-28* endogenous promoter, a *dpy-7* hypodermal promoter (Gilleard et al., 1997), and an *ehn-3A* early somatic gonad promoter (Large and Mathies, 2010). The endogenous *lin-28* promoter drives *lin-28:GFP* expression in neurons and hypodermis, where *lin-28* is known to be expressed (Moss et al., 1997). Using spinning disk microscopy we also detected a low level of GFP expression in Z1 and Z4 cells, which are precursors of somatic gonadal tissues, (Fig S7A,B). *Pdpy-7:lin-28:GFP* was expressed in the hypodermis during the embryonic and L1 stages, and *Pehn-3A::lin-28:GFP* expression was strongest in Z1 and Z4 from the late embryo to L1 stages (Fig S7C,D). *lin-28:GFP* levels decreased from the L2 stage in all three strains, presumably due to repression mediated by the *lin-28* 3’ UTR.

We crossed MosSCI strains with *lin-28(n719);cog-1:GFP* hermaphrodites to assess the relative timing of somatic gonadal and hypodermal *cog-1:GFP* expression. *cog-1:GFP* was expressed in somatic gonadal tissues prior to the vulva in wild-type animals, whereas *cog-1:GFP* expression in the vulva was precocious in the *lin-28(n719)* mutants (Fig 6, 7A,7B). *lin-28* expression via its endogenous promoter *(Plin-28::lin-28:GFP:*lin*-28* 3’UTR) restored the normal relative timing of somatic gonadal and hypodermal *cog-1:GFP* expression in *lin-28(n719)* mutants (Fig 7C). Similarly, in *Pdpy-7::lin-28:GFP;lin-28(n719);cog-1:GFP* animals, *cog-1:GFP* expression in the vulva occurred at the normal time, following somatic gonad expression. Thus, hypodermal expression of *lin-28* efficiently rescues precocious hypodermal development of *lin-28(n719)* (Fig 7D). By contrast, somatic gonadal expression of *lin-28* via the *Pehn-3A::lin-28:GFP* transgene did not rescue the precocious expression of hypodermal *cog-1:GFP* in *lin-28(n719)* mutants (Fig 7E). These results are consistent with cell-intrinsic activity of *lin-28* in the hypodermis to control hypodermal developmental timing.

**Figure 7.**
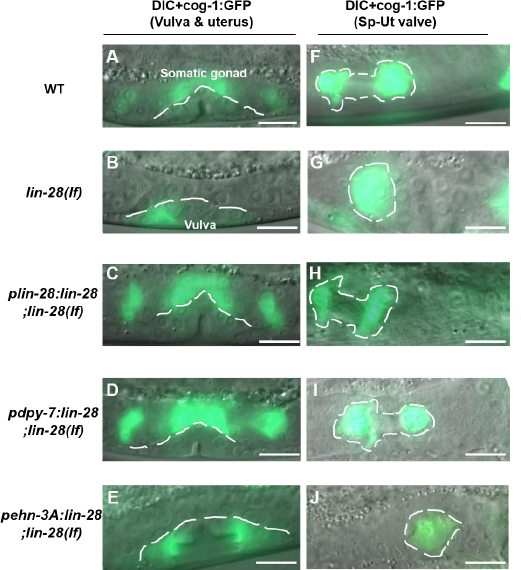
Hypodermal expression of *lin-28* can rescue developmental timing defects and Sp-Ut valve morphogenesis in *lin-28(lf)* mutants. *cog-1.GFP* expression in the somatic gonad and vulva in late 3^rd^ or early 4^th^ larval stage hermaphrodites (A-E), or in the Sp-Ut valve in adult or 4^th^ stage hermaphrodites (F-J), of the wild type (A, F), *lin-28(lf)* (B, G), and *lin-28(f)* with *lin-28* expression driven by various promoters (C-E, H-J). *Pdpy-7* (D, I) is a hypodermal-specific promoter (Gilleard et al., 1997) and *Pehn-3A* (E, J) is an early somatic gonadal-specific promoter (Large and Mathies, 2010). Note: In these experiments, the promoter-driven *lin-28* transgene (C-E, H-J) is also tagged with GFP, but *lin-28:GFP* expression is not detectable at these stages, so all the GFP signal shown here corresponds to *cog-1:GFP. Plin-28::lin-28:GFP;lin-28(n719)* (C) and *Pdpy-7::lin-28:GFP;lin-28(n719)* (D) expresses *cog-1:GFP* in the somatic gonad earlier than in the vulva, as in wild type (A). In contrast, both *lin-28(n719)* mutants (B) and *Pehn-3A::lin-28:GFP;lin-28(n719)* (E) expresses *cog-1:GFP* precociously in the vulva. *Plin-28::lin-28:GFP;lin-28(n719)* (H) and *Pdpy-7::lin-28:GFP;lin-28(n719)* (I) restores the Sp-Ut core dumbbell structure observed in wild type (F). However, the Sp-Ut valve core structure remains as a single lobe shape in both *Pehn-3A::lin-28:GFP;lin-28(n719)* (J) and *lin-28(n719)* (G). These data suggest that hypodermal expression of *lin-28* is sufficient to rescue heterochronic development (A-E) and abnormal Sp-Ut valve core morphogenesis (F-J) in *lin-28(n719)* mutants, but somatic gonadal expression of *lin-28* cannot rescue either defect.

Does hypodermal expression of *lin-28* also rescue the somatic gonadal morphogenesis defects of *lin-28(n719)* hermaphrodites? *lin-28:GFP* driven by the *dpy-7* promoter in *lin-28(n719)* rescued revealed wildtype morphology of the Sp-Ut valve, (Fig 7F, H and I), normal uterine lumen formation(Fig S3I), and normal utse migration (Fig S4M,N). By contrast, somatic gonadal *lin-28* expression did not rescue *lin-28(n719)* somatic gonadal defects (Fig 7G,7J, S3J,S3K, S4O and S4P).

Because the hypodermal expression of *lin-28* suppresses the morphological defects of the somatic gonad in *lin-28(n719)* mutants, we investigated whether it also rescues the fertility defects of these mutants. Expression of either the *lin-28* promoter or *dpy-7* promoter driving *lin-28:GFP* restored the embryo viability of the *lin-28(n719)* mutant to the wild-type level, whereas *ehn-3A* promoter driven *lin-28:GFP;lin-28(n719)* expression did not lead to increased viability compared with *lin-28(n719)* mutants (Fig 8A).

**Figure 8.**
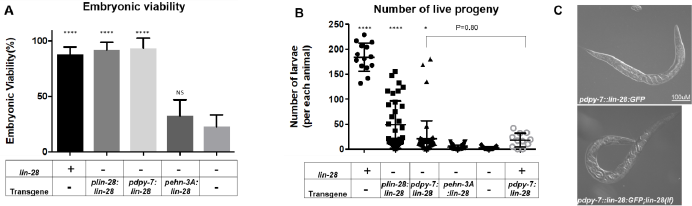
Hypodermal expression of *lin-28* enhances fertility of *lin-28(lf)* mutants. (A) Embryonic viability of *Plin-28::lin-28:GFP;lin-28(n719)* and *Pdpy-7::lin-28:GFP;lin-28(n719)* are restored to a level similar to that in wild type or *lin-2(e1309)* mutants. However, expression of *Pehn-3A::lin-28:GFP* does not enhance the viability of *lin-28(n719)* embryos. (Number of animlals≥15 per each assay; number of independent replicate assays = 4 for *lin-28(n719),* 3 for all other strains.) (B) The number of live larva progeny is increased in *Plin-28::lin-28:GFP;lin-28(n719)* (n=36) and slightly enhanced in *Pdpy-7::lin-28:GFP;lin-28(n719)* (n=53) compared to *lin-28(n719)* mutants (n=25). Progeny numbers for *Pdpy-7:lin-28::GFP* insertional lines without *lin-28(n719)* (n=11) are similar to those of *Pdpy-7::lin-28:GFP;lin-28(n719)* (p value=0.80), suggesting that expression of *Pdpy-7:lin-28::GFP* induces fertility defects regardless of *lin-28(n719).* Progeny numbers for *Pehn-3A::lin-28:GFP;lin-28(n719)* (n=40) are similar to those of *lin-28(n719)* mutants. (A,B: Unpaired t-test compared to *lin-28(lf),* NS; not significant, *p<0.05,****p<0.0001) (C) Embryos of *Pdpy-7::lin-28:GFP* (upper panel) and *Pdpy-7::lin-28:GFP;lin-28(n719)* (lower panel) are trapped inside adult hermaphrodites, indicating that defective egg laying is a cause of the reduced fertility in these animals. (Scale bar = 100 µm)

The total number of live progeny was greater for the *Pdpy-7::lin-28:GFP;lin-28(n719)* strain that for the *lin-28(n719)* mutants. However, only ~15% of animals had numbers of progeny comparable with wild-type values, and the other ~85% of animals produced less than 20 progeny at 25°C (Fig 8B). In addition, *Pdpy-7::lin-28:GFP* insertional strains in the wild-type background led to a similar number of progeny, indicating that the expression of *Pdpy-7::lin-28:GFP* intrinsically induces the fertility defects. We found that egg-laying is defective for *Pdpy-7::lin-28:GFP;lin-28(n719)* and *Pdpy-7::lin-28:GFP* hermaphrodites (Fig 8C). Vulval functions for egg-laying in those strains were intact though, because the animals were able to lay eggs occasionally. In contrast, somatic gonadal promoter-driven lin-28:GFP expression did not affect the total number of progeny in *lin-28(n719)* animals (Fig 8B).

## Discussion

*C. elegans lin-28(lf)* hermaphrodites exhibit a dramatic reduction in fertility, to a degree in excess of what would be expected as a simple consequence of their vulval morphogenesis defects. In principle, it was possible that *lin-28* could promote fertility entirely via a germline-specific activity, analogous to the demonstrated role of mammalian Lin28 in promoting pluripotency and stem cell proliferation (Asara et al., 2016; Yu et al., 2007). Indeed, *lin-28* has been reported to regulate the germ cell pool size in mice and in *C. elegans* hermaphrodites (Shinoda et al., 2013; Wang et al., 2017). However, the reported effects on germ cell pool size in *C. elegans,* after germline knockdown of *lin-28,* did not include substantially reduced fertility (Wang et al., 2017). We performed tissue-specific RNAi experiments using *rrf-1(lf)* and *ppw-1(lf),* and observed that somatic knockdown of *lin-28* caused greater reduction of fertility than did germline knockdown of *lin-28* (Fig S8). This suggests that the dramatically reduced fertility of *lin-28(lf)* hermaphrodites is caused by fertility-promoting activities of *lin-28* outside of the germline in *C. elegans,* that is, within somatic cell lineages of the gonad and/or other tissues. Other examples in which *lin-28* controls the development of reproductive tissues other than germ cells have been reported in flies and mice. The *Drosophila* egg chamber is fused abnormally early in *lin-28* null mutants, and the development of mice vaginal openings is delayed in *lin-28a* transgenic mice (Stratoulias et al., 2014; Zhu et al., 2010).

Here we investigated the somatic function of *lin-28* in promoting *C. elegans* hermaphrodite fertility. Our results show that *lin-28* is required for the completion of certain somatic gonadal morphogenetic events, specifically, the enlargement of the uterine lumen, the migration of utse nuclei, and extension of the Sp-Ut valve core, and that these somatic gonadal morphogenesis defects likely underlie the reduced fertility of *lin-28(lf)* hermaphrodites. In particular, the abnormal Sp-Ut valve of *lin-28(lf)* animals has a potent impact on fertility by preventing fertilized embryos from entering the uterus, thereby inhibiting ovulation and resulting in reduced embryo production.

We found that egg shell integrity is also compromised in *lin-28(lf)* mutants, which negatively affects embryonic viability, similarly to other egg shell defective mutants (Johnston et al., 2006; Johnston et al., 2010; Maruyama et al., 2007). Moreover, we found that another spermathecal exit mutant *fln-1(lf)* has egg shell-defective phenotypes similar to *lin-28(lf)* (Fig 4C), suggesting a causal link between spermathecal retention and egg shell integrity. We speculate that physical damage to the egg shells may occur when the embryos are trapped in the spermathecae. For example, the pressure induced by oocytes in the proximal gonad that are readily ovulated might serve as one factor. In addition, *C. elegans* spermathecae might be a harsh environment for the maintenance of egg shell integrity over a long time. It is reported that the pH of spermathecal fluid is high (pH=8.6) in queen bees, *Apis mellifera* (Gessner and Gessner, 1976). Interestingly, deficiency of *cbd-1,* an essential component of *C. elegans* egg shell, leads to pinched off embryos reflecting incomplete spermathecal exit (Johnston et al., 2010). This suggests that damage to the egg shell of embryos lingering too long in the spermatheca could further aggravate an underlying spermathecal exit defect. Further research is needed to clarify the relationship between spermathecal exit and egg shell integrity.

Our results point to a close coupling between the hypodermal and somatic gonadal phenotypes in *lin-28(lf)* mutants. Firstly, we found that a very similar configuration of the known heterochronic genes mediates the effects of *lin-28(lf)* on hypodermal developmental timing and on somatic gonadal morphogenesis (Fig 5E). The only exception could be *lin-41;* we could not find evidence that *lin-41,* a downstream target of *let-7* for hypodermal developmental timing (Slack et al., 2000), is involved in Sp-Ut valve morphogenesis. The lin-41(RNAi) animals displayed a superficially normal dumbbell-shaped Sp-Ut valve, albeit somewhat smaller than the wild-type valve (Fig S6C). This result may reflect either an insufficient knockdown of *lin-41* by RNAi in these experiments, or that *lin-41* does not participate in somatic gonad development. In the latter case, *hbl-1* would appear to function as a main downstream target of *let-7* in somatic gonad morphogenesis.

Our second finding indicating a linkage between *lin-28(lf)* hypodermal and somatic gonadal phenotypes, is that post-dauer development suppresses both the hypodermal and the somatic gonad developmental defects of *lin-28(lf)* hermaphrodites. This latter result in particular accentuates that the somatic gonadal defects of *lin-28(lf)* do not reflect a direct role of *lin-28* in somatic gonadal development, *per se;* rather, it seems that somatic gonadal morphogenesis fails in *lin-28* mutants as an indirect consequence of precocious development. Consistent with this idea, another precocious mutant *lin-14(lf)* showed similar Sp-Ut valve defects to *lin-28(lf)* mutants (Fig S6D). L1-specific cell fates were skipped in the hypodermis of *lin-14(lf)* mutants (Ambros and Horvitz, 1984), and thus, these mutants go through only three larval stages, just as observed for *lin-28(lf)* mutants. However, we can also interpret this result to mean that a reduced *lin-28* level in the *lin-14(lf)* mutant causes phenotypes similar to those of the *lin-28(lf)* mutants, because *lin-14* positively regulates *lin-28* (Pepper et al., 2004; Seggerson et al., 2002).

Importantly, we did not find evidence for precocious development of somatic gonadal events in *lin-28(lf).* In particular, we observed that the timing of the onset of expression of certain fluorescent markers of L3 and L4 somatic gonadal developmental events was not altered in *lin-28(lf),* even though subsequent morphogenesis failed. Assuming that the somatic gonad and the hypodermis each has its own developmental clock, it would appear that the hypodermal developmental clock of *lin-28(lf)* mutants is accelerated, while the somatic gonadal developmental clock runs normally, resulting in discoordination of developmental timing between the two tissues during the L3 and L4 stages (Fig 6F). We propose that it is this temporal discord between the accelerated hypodermis, and the normally-scheduled somatic gonad, that results in failure of somatic gonadal morphogenesis.

The apparent absence of precocious development of the somatic gonad of *lin-28(lf)* animals suggests that *lin-28* may affect somatic gonadal morphogenesis cell non-autonomously, by controlling one or more signals from the hypodermis to the somatic gonad. In strong support for this model, we found that expression of *lin-28* specifically in the hypodermis could rescue the somatic gonadal developmental defects of *lin-28(lf)* mutants. Conversely, expression of *lin-28* specifically in the somatic gonadal precursor lineage did not rescue any *lin-28(lf)* phenotypes. Based on these observations, we propose that the principle function of *lin-28* with regards to somatic gonadal development is to act within the hypodermis to specify a schedule of hypodermal events that is properly coordinated with a corresponding schedule of somatic gonadal developmental events. Accordingly, normal somatic gonadal morphogenesis is proposed to require a coordinated agenda of signaling between the hypodermis and the somatic gonad during the L3 and/or L4 stages.

Then, by what mechanisms could hypodermal activity of *lin-28* regulate the development of a different tissue? There are precedents in *C. elegans* for cell non-autonomous developmental signals originating from the hypodermis. A recent study showed that heterochronic genes acting in the hypodermis can modulate mTOR signaling in the intestine (Dowen et al., 2016). This signaling requires the mTORC2 complex specifically and its downstream factors including *rict-1/rictor, sinh-1/sin, and sgk-1/sgk1* in the intestine. However, it is unlikely that somatic gonad development is related to mTORC2 signaling, because we found that *rict-1* (RNAi) in *lin-28(lf)* mutants did not induce or suppress Sp-Ut valve defects (data not shown). In another example, it has been reported that migration of the hermaphrodite-specific neurons and arborization of sensory neurons are regulated by hypodermal expression of the microRNA *mir-79* and MNR-1/menorin, respectively (Pedersen et al., 2013; Salzberg et al., 2013). Hypodermal glycosylated cell surface molecules or signaling cell adhesion molecules are key downstream factors for neuronal morphogenesis in each case, implying this nonautonomous signaling may require physical contact with other tissues.

Indeed, seam cells in the hypodermis become connected to the utse (uterine seam cell) of *C. elegans* hermaphrodites. The connection between these two tissues is thought to be formed during the mid to late L4 stages in wild-type animals (Newman et al., 1996). The precocious hypodermal maturation in *lin-28(lf)* animals might cause this connection to be formed aberrantly, or not at all. Alternatively, the reduced number of seam cells in *lin-28(lf)* mutants may alter positioning of seam cells in the hypodermis, disrupting the normal connection between seam cells and utse. However, there is evidence that utse defects may not necessarily be associated with Sp-Ut abnormality. *lin-29* expression in the anchor cell induces signals for utse precursor cells to adopt utse fates, and *lin-29(lf)* mutants do not form a proper utse (Newman et al., 2000). However, loss of *lin-29* does not cause any detectable abnormality in Sp-Ut valve core morphology (Fig 5A). Moreover, it has been reported that anchor cell invasion genes *(fos-1, mig-10, egl-43, cdh-3, and zmp-1)* are involved in utse development (Ghosh and Sternberg, 2014), but we did not observe any abnormality in the Sp-Ut valve upon RNAi knockdown of these genes (data not shown). Nevertheless, it will be interesting to examine whether the physical connection between the hypodermal seam and gonadal utse is formed properly in *lin-28(lf)* hermaphrodites.

It is striking that *lin-28(lf)* mutants exhibit coordinate defects in the final stages of morphogenesis in at least three distinct somatic gonadal structures: extension of the Sp-Ut valve core, positioning of the uterine seam cell nuclei, and expansion of the lumen of the uterus. Since all three of these defects are highly penetrant in *lin-28(lf),* and are coordinately rescued by appropriate lin-28-expressing transgenes, we were not able to determine if these defects are expressed independently, or whether for example, one of them is the primary defect that is linked to hypodermal developmental timing, and the other defects are secondarily precipitated the first. Further research is needed to identify any cause-effect relationships between utse migration, Sp-Ut valve morphogenesis, and uterus lumen formation.

Overall, our studies of *lin-28* in the context of *C. elegans* reproductive system development provides an informative model for exploring fundamental principles of multicellular development, including how the generation of organized cellular complexity requires precise temporal coordination of events across interacting tissues. Our findings exemplify how a cell-intrinsic developmental timing program can be required not only cell-autonomously to specify temporal cell fate, but also to control cell-nonautonomous signaling that is critical for proper development of interacting tissues. Our identification of cell-nonautonomous hypodermis-to-gonad developmental signaling controlled by *lin-28* and the heterochronic pathway should set the stage for future studies addressing the identity and potential evolutionary conservation of the molecular components of the downstream signal(s).

## Material and Methods

### Culture of *C. elegans* strains

*C. elegans* wild type (Strain N2) and mutant strains (listed in Supplementary Table 1) were grown and maintained (at 25°C unless otherwise noted) on nematode growth media (NGM) agar plates seeded with *E. coli* (strain HB101). A list of genotyping primers for allele confirmation can be found in Supplementary Table 2. Synchronized populations of larvae at defined developmental stages were obtained as previously described (Stiernagle, 2006). Briefly, embryos were collected using sodium hypochlorite and 5N NaOH, washed with M9 buffer, and incubated in M9 buffer overnight at 20°C, placed on NGM plates seeded with HB101, incubated for defined lengths of time at 20°C or 25°C, and developmental stage was assessed by DIC microscopy of a sample of worms from the population (Byerly et al., 1976).

### Microscopy

For DIC and fluorescence microscopy, worms were anesthetized with 0.2mM levamisol and mounted on 2% agarose pads. All images except the following were obtained with a ZEISS Axiocam 503 mono: Supplementary Figure 2 (Leica DM 5500Q confocal microscopy), Supplementary Figure 7 (3i (Intelligent imaging Innovations) Everest spinning disk confocal microscopy). The supplementary videos were taken with a Zeiss Axioplan2. For gonad DAPI staining, hermaphrodites were cut with a syringe needle and the extruded gonads were fixed with 95% ethanol. After washing twice with M9, the dissected gonads were incubated with 4’6’-diamidino-2-phenylindole solution (100ng/ml) for 10min in a humidified chamber and washed again with M9 (Modified from (Shaham, 2005)).

### RNAi

RNAi by feeding worms with *E. coli* expressing double-stranded RNA was conducted as previously described (Conte et al., 2015). HT115 bacterial RNAi strains *(lin-28, hbl-1, lin-29, lin-46, rict-1, fos-1, mig-10, egl-43, cdh-3, zmp-1)* and an empty vector strain (L4440) from the Ahringer library were used (Kamath and Ahringer, 2003).

### Analysis of fertility

Individual young adult hermaphrodites were placed, one per plate, on a NGM plates seeded with HB101, and the number of live progeny from each hermaphrodite was counted 3~4 days later. To determine embryo viability, gravid adult hermaphrodites were dissected with a syringe, the released embryos were collected, and transferred to NGM plates seeded with HB101. The total number of embryos was counted immediately, and after 36hrs incubation at 25°C, the number of live animals was counted. Viability was calculated as (Number of live animals / Total number of embryos seeded) X 100%.

### Egg shell integrity

Egg shell permeability was asessed using FM 4-64 dye (Sigma,T13320), as described (Johnston et al., 2006). Briefly, embryos were dissected from gravid hermaphrodites in 150mM KCl with 30uM of FM4-64, and the proportion of embryos infiltrated by FM4-64 was measured using fluorescence microscopy.

### Construction of plasmids

To generate transgenic strains containing tissue specific promoters driving *lin-28:GFP,* we removed the sequence between the first exon and second exon (I:8409341-I:8410415) of *lin-28a* to prevent the sequence from serving as an endogenous promoter (Moss et al., 1997). GFP sequences were adapted from XW12 (Wei et al., 2012) and were fused in frame to the carboxy terminus of *lin-28* coding sequence. The primers used for the overlapping PCRs for these procedures are listed in Supplementary Table 2. We used Gateway^®^ Technology (Invitrogen, cat 12535-019) to construct transgenic vectors. *lin-28:GFP* was cloned into the gateway entry vector pDONR P2P3 by BP reaction. Also, the promoter regions of *enh-3A, lin-28,* and *dpy-7* were cloned into pDONR P4P1r by BP reaction. *lin-28* 3’UTR was cloned into pDONR P2rP3. LR reactions of these three entry vectors with pCJF150 yielded final vectors containing the following transgenes for injection: pSW40(plin-28), pSW42(pehn-3A), and pSW43(pdpy-7). The sequences for all primers used in this procedure can be found in Supplementary Table 2 and the maps of pSW40, pSW42, and pSW43 can be found in Supplementary Figure 9.

### Generation of MosSCI transgenic lines

MosSCI single-copy insertions (into the *ttTi5605* Mos1 allele, near the center of chromosome II) were obtained using the protocol previously described on the “wormbuilder” website (http://www.wormbuilder.org/). For each plasmid construct, MosSCI transformation was generally conducted using the direct injection approach (Frøkjær-Jensen et al., 2012), and also using the approach employing extrachromosomal array intermediates and ivermectin selection for insertion (Shirayama et al., 2012). To prepare plasmids for injection, the following plasmids were purified using a midiprep kit (Qiagen, Cat.12143): pSW40, pSW42, PSW43, pCFJ601 (Peft-3::transposase), pMA122 (Phsp::peel-1), pGH8 (Prab-3::mCherry), pCFJ90 (Pmyo-2::mCherry), pCFJ104(Pmyo-3::mCherry), pJL44(Phsp-6.48::MosTase::glh-2 3’UTR), pCCM416(Pmyo-2::avr-15), and *pRF4(rol-6(su1006)).* For the direct injection method, injection mixtures consisted of pCFJ601 (50ng/ul), pMA122(10ng/ul), pGH8(10ng/ul), pCFJ90(2.5ng/ul), pCFJ104(5ng/ul), and one of the transgene-containing plasmids (pSW40, pSW42, or pSW43(25ng/ul); the mixture was injected into EG4322 hermaphrodites and injected animals were placed singly onto NGM plates seeded with HB101. Following 7~10 days of incubation at 25°C, cultures were heat-shocked (35°C, 1hr) to kill any worms with an extrachromosomal array and surviving animals were cloned. After allowing them to produce progeny, worms are genotyped to identify single copy transgene inserted strains. For the approach using ivermectin selection, we injected a mixture of pJL44(50ng/ul), pCCM416(50ng/ul), and pRF4(50ng/ul) with one of pSW40, pSW42, or pSW43(15ng/ul) into WM186 hermaphrodites. After allowing the progeny of injected animals to grow for multiple generations, they were heat-shocked (35°C, 1hr) to induce heat shock promoter driven transposase expression from extra chromosomal arrays, and single single-copy transgene inserted strains were selected by invermectin resistance (10ng/ml) against extrachromosomal arrays. We obtained VT3702 by the direct injection method, and VT3486 and VT3392 by the extrachromosomal array intermediate method. We crossed those strains to VT2929 to obtain VT3703, VT3517 and VT3516 (Supplementary Table 1).

## Acknowledgement

We would like to thank all Ambros Lab members, Anna Zinovyeva (Kansas State University), Katherine McJunkin (NIH), and Yoshiki Andachi (National Institute of Genetics) for helpful discussions and critical comments about this project. We also would like to thank Erin Cram (Northeastern University) and Anna Allen (Howard University) for sharing strains, Kang Shen (Stanford University) and Sean Ryder (University of Massachusetts Medical School) for sharing plasmids and Amy Walker, Craig Mello, and Michael Gorzyca (University of Massachusetts Medical School) for help to use their microscopes.

## Funding

### National Institutes of Health (R01GM104904-04)

Victor Ambros

### National Institutes of Health (R01GM034028-32)

Victor Ambros

The funders had no role in study design, data collection and interpretation, or the decision to submit the work for publication.

## References

Abbott, A. L., Alvarez-Saavedra, E., Miska, E. a, Lau, N. C., Bartel, D. P., Horvitz, H. R. and Ambros, V. (2005). The let-7 MicroRNA family members mir-48, mir-84, and mir-241 function together to regulate developmental timing in Caenorhabditis elegans. Dev. cell 9, 403–14.

Abrahante, J. E., Daul, A. L., Li, M., Volk, M. L., Tennessen, J. M., Miller, E. A. and Rougvie, A. E. (2003). The Caenorhabditis elegans hunchback-like Gene lin-57/hbl-1 Controls Developmental Time and Is Regulated by MicroRNAs. Dev. Cell 4, 625–637.

Ambros, V. (2011). MicroRNAs and developmental timing. Curr. Opin. Genet. & Dev. 21, 511–7.

Ambros, V. and Horvitz, H. (1984). Heterochronic mutants of the nematode Caenorhabditis elegans. Science 226, 409–416.

Ambros, V. and Horvitz, H. R. (1987). The lin-14 locus of Caenorhabditis elegans controls the time of expression of specific postembryonic developmental events. Genes & Dev. 1, 398–414.

Asara, J. M., Trapnell, C., Mikkelsen, T., Xia, Q., Loh, Y. H., Loh, Y. H., Teitell, M. A., Shinoda, G., Xie, W., Cahan, P., et al. (2016). LIN28 Regulates Stem Cell Metabolism and Conversion to Primed Pluripotency. Cell Stem Cell 19, 66–80.

Byerly, L., Cassada, R. C. and Russell, R. L. (1976). The life cycle of the nematode Caenorhabditis elegans. I. Wild-type growth and reproduction. Dev. Biol. 51, 23–33.

Chang, W., Tilmann, C., Thoemke, K., Markussen, F.-H., Mathies, L. D., Kimble, J. and Zarkower, D. (2004). A forkhead protein controls sexual identity of the C. elegans male somatic gonad. Development. 131, 1425–36.

Conte, D., MacNeil, L. T., Walhout, A. J. M. and Mello, C. C. (2015). RNA Interference in Caenorhabditis elegans. Curr. Protoc. Mol. Biol. 109, 26.3.1–26.330.

Dowen, R. H., Breen, P. C., Tullius, T., Conery, A. L. and Ruvkun, G. (2016). A microRNA program in the C. elegans hypodermis couples to intestinal mTORC2/PQM-1 signaling to modulate fat transport. Genes Dev 30, 1515–1528.

Euling, S. and Ambros, V. (1996). Heterochronic genes control cell cycle progress and developmental competence of C-elegans vulva precursor cells. Cell 84, 667–676.

Euling, S. and Ambros VR (1996). Reversal of cell fate determination in Caenorhabditis elegans vulval development. Development 122, 2507–15.

Frokjær-Jensen, C., Davis, M. W., Ailion, M. and Jorgensen, E. M. (2012). Improved Mos1-mediated transgenesis in C. elegans. Nat. methods 9, 117–118.

Gessner, B. and Gessner, K. (1976). Inorganic ions in spermathecal fluid and their transport across the spermathecal membrane of the queen bee, Apis mellifera. J. Insect Physiol. 22, 1469–1474.

Ghosh, S. and Sternberg, P. W. (2014). Spatial and molecular cues for cell outgrowth during C. elegans uterine development. Dev. Biol. 396, 121–135.

Gilleard, J. S., Barry, J. D. and Johnstone, I. L. (1997). cis regulatory requirements for hypodermal cell-specific expression of the Caenorhabditis elegans cuticle collagen gene dpy-7. Mol. Cell. Biol. 17, 2301–2311.

Gissendanner, C. R., Kelley, K., Nguyen, T. Q., Hoener, M. C., Sluder, A. E. and Maina, C. V. (2008). The Caenorhabditis elegans NR4A nuclear receptor is required for spermatheca morphogenesis. Dev. Biol. 313, 767–786.

Hoskins, R., Hajnal, a F., Harp, S. a and Kim, S. K. (1996). The C. elegans vulval induction gene lin-2 encodes a member of the MAGUK family of cell junction proteins. Development. 122, 97–111.

Iwasaki, K., James, M., Francis, R. and Schedl, T. (1996). emo-1, a Caenorhabditis elegans Sec61 p r Homologue, Is Required for Oocyte Development and Ovulation. J. cell Biol. 134, 699–714.

Johnston, W. L., Krizus, A. and Dennis, J. W. (2006). The eggshell is required for meiotic fidelity, polar-body extrusion and polarization of the C. elegans embryo. BMC Biol. 4, 35.

Johnston, W. L., Krizus, A. and Dennis, J. W. (2010). Eggshell chitin and chitin-interacting proteins prevent polyspermy in C. elegans. Curr. Biol.: CB 20, 1932–7.

Kariya, K.-I., Bui, Y. K., Gao, X., Sternberg, P. W. and Kataoka, T. (2004). Phospholipase Cepsilon regulates ovulation in Caenorhabditis elegans. Dev. Biol. 274, 201–10.

Keyte, A. L. and Smith, K. K. (2012). Heterochrony and developmental timing mechanisms: Changing ontogenies in evolution. Semin. Cell & Dev. Biol. 34, 99–107.

Kimble, J. and Hirsh, D. (1979). The postembryonic cell lineages of the hermaphrodite and male gonads in Caenorhabditis elegans. Dev. Biol. 70, 396–417.

Klingenberg, C. P. (1998). Heterochrony and allometry: the analysis of evolutionary change in ontogeny. Biol. Rev. Camb. Philos. Soc. 73, 79–123.

Kovacevic, I. and Cram, E. J. (2010). FLN-1/filamin is required for maintenance of actin and exit of fertilized oocytes from the spermatheca in C. elegans. Dev. Biol. 347, 247–57.

Large, E. E. and Mathies, L. D. (2010). hunchback and Ikaros-like zinc finger genes control reproductive system development in Caenorhabditis elegans. Dev. Biol. 339, 51–64.

Lee, R. C., Feinbaum, R. L. and Ambros, V. (1993). the C. elegans\rheterochronic gene lin-4 encodes small RNAs with antisense\rcomplementarity to lin-14. Cell 75: 843–85, 843-854.

Liu, Z. and Ambros, V. (1991). Alternative temporal control systems for hypodermal cell differentiation in Caenorhabditis elegans. Nature 350, 162–165.

Maruyama, R., Velarde, N. V., Klancer, R., Gordon, S., Kadandale, P., Parry, J. M., Hang, J. S., Rubin, J., Stewart-Michaelis, A., Schweinsberg, P., et al. (2007). EGG-3 regulates cell-surface and cortex rearrangements during egg activation in Caenorhabditis elegans. Curr. Biol.: CB 17, 1555–60.

Mayr, F., Schütz, A., Döge, N. and Heinemann, U. (2012). The Lin28 cold-shock domain remodels pre-let-7 microRNA. Nucleic Acids Res. 40, 7492–7506.

McCarter, J., Bartlett, B., Dang, T. and Schedl, T. (1999). On the control of oocyte meiotic maturation and ovulation in Caenorhabditis elegans. Dev. Biol. 205, 111–28.

Moss, E. G., Lee, R. C. and Ambros, V. (1997). The Cold Shock Domain Protein LIN-28 Controls Developmental Timing in C. elegans and Is Regulated by the lin-4 RNA. Cell 88, 637–646.

Newman, A. P. and Sternberg, P. W. (1996). Coordinated morphogenesis of epithelia during development of the Caenorhabditis elegans uterine-vulval connection. Proc. Natl. Acad. Sci. United States Am. 93, 9329–9333.

Newman, A. P., White, J. G. and Sternberg, P. W. (1996). Morphogenesis of the C. elegans hermaphrodite uterus. Development. 122, 3617–3626.

Newman, A. P., Inoue, T., Wang, M. and Sternberg, P. W. (2000). The Caenorhabditis elegans heterochronic gene lin-29 coordinates the vulval-uterine-epidermal connections. Curr. Biol. 10, 1479–1488.

Palmer, R. E., Inoue, T., Sherwood, D. R., Jiang, L. I. and Sternberg, P. W. (2002). Caenorhabditis elegans cog-1 Locus Encodes GTX/Nkx6.1 Homeodomain Proteins and Regulates Multiple Aspects of Reproductive System Development. Dev. Biol. 252, 202–213.

Pedersen, M. E., Snieckute, G., Kagias, K., Nehammer, C., Multhaupt, H. A. B., Couchman, J. R. and Pocock, R. (2013). An epidermal microRNA regulates neuronal migration through control of the cellular glycosylation state. Science. 341, 1404–1408.

Pepper, A. S.-R., McCane, J. E., Kemper, K., Yeung, D. A., Lee, R. C., Ambros, V. and Moss, E. G. (2004). The C. elegans heterochronic gene lin-46 affects developmental timing at two larval stages and encodes a relative of the scaffolding protein gephyrin. Developement. 131, 2049–59.

Piskounova, E., Polytarchou, C., Thornton, J. E., LaPierre, R. J., Pothoulakis, C., Hagan, J. P., Iliopoulos, D. and Gregory, R. I. (2011). Lin28A and Lin28B inhibit let-7 microRNA biogenesis by distinct mechanisms. Cell 147, 1066–79.

Reinhart, B. J., Slack, F. J., Basson, M., Pasquinelli, a E., Bettinger, J. C., Rougvie, a E., Horvitz, H. R. and Ruvkun, G. (2000). The 21-nucleotide let-7 RNA regulates developmental timing in Caenorhabditis elegans. Nature 403, 901–6.

Resnick, T. D., McCulloch, K. a and Rougvie, A. E. (2010). miRNAs give worms the time of their lives: small RNAs and temporal control in Caenorhabditis elegans. Dev. Dyn.: Off. Publ. Am. Assoc. Anat. 239, 1477–89.

Salzberg, Y., Díaz-Balzac, C. A., Ramirez-Suarez, N. J., Attreed, M., Tecle, E., Desbois, M., Kaprielian, Z. and Bülow, H. E. (2013). Skin-Derived Cues Control Arborization of Sensory Dendrites in Caenorhabditis elegans. Cell 155, 10.1016/j.cell.2013.08.058.

Seggerson, K., Tang, L. and Moss, E. G. (2002). Two genetic circuits repress the Caenorhabditis elegans heterochronic gene lin-28 after translation initiation. Dev. Biol. 243, 215–25.

Shaham, S. (2005). Methods in cell biology. WormBook.

Shinoda, G., de Soysa, T. Y., Seligson, M. T., Yabuuchi, A., Fujiwara, Y., Yi Huang, P., Hagan, J. P., Gregory, R. I., Moss, E. G. and Daley, G. Q. (2013). Lin28a Regulates Germ Cell Pool Size and Fertility. Stem cells 2013,.

Slack, F. J., Basson, M., Liu, Z., Ambros, V., Horvitz, H. R. and Ruvkun, G. (2000). The lin-41 RBCC Gene Acts in the C. elegans Heterochronic Pathway between the let-7 Regulatory RNA and the LIN-29 Transcription Factor. Mol. Cell 5, 659–669.

Shirayama, M., Seth, M., Lee, H.-C., Gu, W., Ishidate, T., Conte, D. and Mello, C. C. (2012). piRNAs initiate an epigenetic memory of nonself RNA in the C. elegans germline. Cell 150, 65–77.

Stratoulias, V., Heino, T. I. and Michon, F. (2014). Lin-28 regulates oogenesis and muscle formation in Drosophila melanogaster. PLoS ONE 9, 1–14.

Stiernagle, T. (2006). Maintenance of C. elegans (February 11, 2006), WormBook, ed. The C. elegans Research Community, WormBook, doi/10.1895/wormbook. 1.101. 1.

Tompkins, R. (1978). Genie control of axolotl metamorphosis. Integr. Comp. Biol. 18, 313–319.

Vadla, B., Kemper, K., Alaimo, J., Heine, C. and Moss, E. G. (2012). lin-28 Controls the Succession of Cell Fate Choices via Two Distinct Activities. PLoS Genet. 8, e1002588.

Van Wynsberghe, P. M., Kai, Z. S., Massirer, K. B., Burton, V. H., Yeo, G. W. and Pasquinelli, A. E. (2011). LIN-28 co-transcriptionally binds primary let-7 to regulate miRNA maturation in C. elegans. Nat Struct Mol Biol 18, 302–308.

Viswanathan, S. R., Daley, G. Q. and Gregory, R. I. (2008). Selective blockade of microRNA processing by Lin28. Science 320, 97–100.

Wang, D., Hou, L., Nakamura, S., Su, M., Li, F., Chen, W., Yan, Y., Green, C. D., Chen, D., Zhang, H., et al. (2017). LIN-28 balances longevity and germline stem cell number in Caenorhabditis elegans through let-7/AKT/DAF-16 axis. Aging Cell 16, 113–124.

Wei, X., Potter, C. J., Luo, L. and Shen, K. (2012). Controlling gene expression with the Q repressible binary expression system in Caenorhabditis elegans. Nat. methods 9, 391–5.

Yu, J., Vodyanik, M. A., Smuga-Otto, K., Antosiewicz-Bourget, J., Frane, J. L., Tian, S., Nie, J., Jonsdottir, G. A., Ruotti, V., Stewart, R., et al. (2007). Induced Pluripotent Stem Cell Lines Derived from Human Somatic Cells. Science 318, 1917–1920.

Zhang, Y., Foster, J. M., Nelson, L. S., Ma, D. and Carlow, C. K. S. (2005). The chitin synthase genes chs-1 and chs-2 are essential for C. elegans development and responsible for chitin deposition in the eggshell and pharynx, respectively. Dev. Biol. 285, 330–9.

Zhu, H., Shah, S., Shyh-Chang, N., Shinoda, G., Einhorn, W. S., Viswanathan, S. R., Takeuchi, A., Grasemann, C., Rinn, J. L., Lopez, M. F., et al. (2010). Lin28a transgenic mice manifest size and puberty phenotypes identified in human genetic association studies. Nat. Genet. 42, 626–30.

Zhu, H., Ng, S. C., Segr, A. V., Shinoda, G., Shah, S. P., Einhorn, W. S., Takeuchi, A., Engreitz, J. M., Hagan, J. P., Kharas, M. G., et al. (2011). The Lin28/let-7 axis regulates glucose metabolism. Cell 147, 81–94.

